# Gcm alleviates the inflammatory phenotype induced by Toll activation in *Drosophila* hemocytes

**DOI:** 10.1101/2023.09.21.558811

**Authors:** Wael Bazzi, Sara Monticelli, Claude Delaporte, Céline Riet, Angela Giangrande, Pierre B. Cattenoz

## Abstract

Hemocytes, the myeloid-like immune cells of *Drosophila*, fulfil a variety of functions that are not completely understood, ranging from phagocytosis to transduction of inflammatory signals. We here show that downregulating the hemocyte-specific Glide/Gcm transcription factor enhances the inflammatory response to the constitutive activation of the Toll pathway. This correlates with lower levels of glutathione S transferase, suggesting an implication of Glide/Gcm in ROS signaling and calling for a widespread anti-inflammatory potential of Glide/Gcm. We show the expression of neurotransmitter receptors in hemocytes and that Toll activation affects their expressions, disclosing a novel aspect of the inflammatory response mediated by neurotransmitters. Finally, we provide evidence for acetylcholine receptor nAchRalpha6 regulating hemocyte proliferation. Altogether, this study provides new insights on the molecular pathways involved in the inflammatory response.

## Introduction

Inflammation is the first response of the organism to pathogenic cues and tissue damages. It allows the removal of the infectious agent and induces the healing process. Prolonged or chronic activation of the inflammatory response is highly detrimental for the organism and constitutes a major aggravating factor in the etiology of many diseases ranging from cancers to neurodegenerative disorders (1-3). Thus, the coordination of the inflammatory response requires robust regulatory mechanisms to prevent its adverse effects.

The inflammatory response is well conserved across evolution and the *Drosophila* model has been instrumental for the identification of the molecular mechanisms underlying innate immunity (4). Two major signalling pathways transducing the inflammatory response are the Toll and the Janus kinase/signal transducer and activator of transcription (Jak/Stat) pathways. Microbial particles activate the Toll receptor, which promotes the degradation of the NfκB inhibitor Cactus (i.e. IκB in mammals), hence allowing the nuclear translocation of the NfκB transcription factors Dorsal (Dl) and Dorsal-related immunity factor (Dif) and the transcription of effector genes (5, 6). The Jak/Stat pathway is activated in response to cytokine signalling. Following the neutralization of the pathogen, the restoration of homeostasis requires the inhibition off of the inflammatory pathways, which depends on potent negative autoregulatory loops where each pathway activates its own inhibitors (7, 8).

*Drosophila* hemocytes are myeloid-like cells that respond to inflammatory challenges. Like in vertebrates, they are produced by two hematopoietic waves occurring in different anlagens and times. The 1^st^ wave hemocytes originate from the procephalic mesoderm of the embryo and circulate in the larval hemolymph or reside between the muscles and the cuticle (*i*.*e*. sessile pockets, dorsal stripes) (**Figure 1A**) (9, 10). The 2^nd^ wave occurs in the larval lymph gland, which hystolizes and releases a second pool of hemocytes after puparium formation (or already in the larva, upon immune challenge) (9). The Toll and the Jak/Stat pathways activate the hemocytes originating from the two hematopoietic waves, leading to their differentiation into lamellocytes, large cells that encapsulate pathogens too big to be phagocytosed (11, 12).

**Figure 1:**
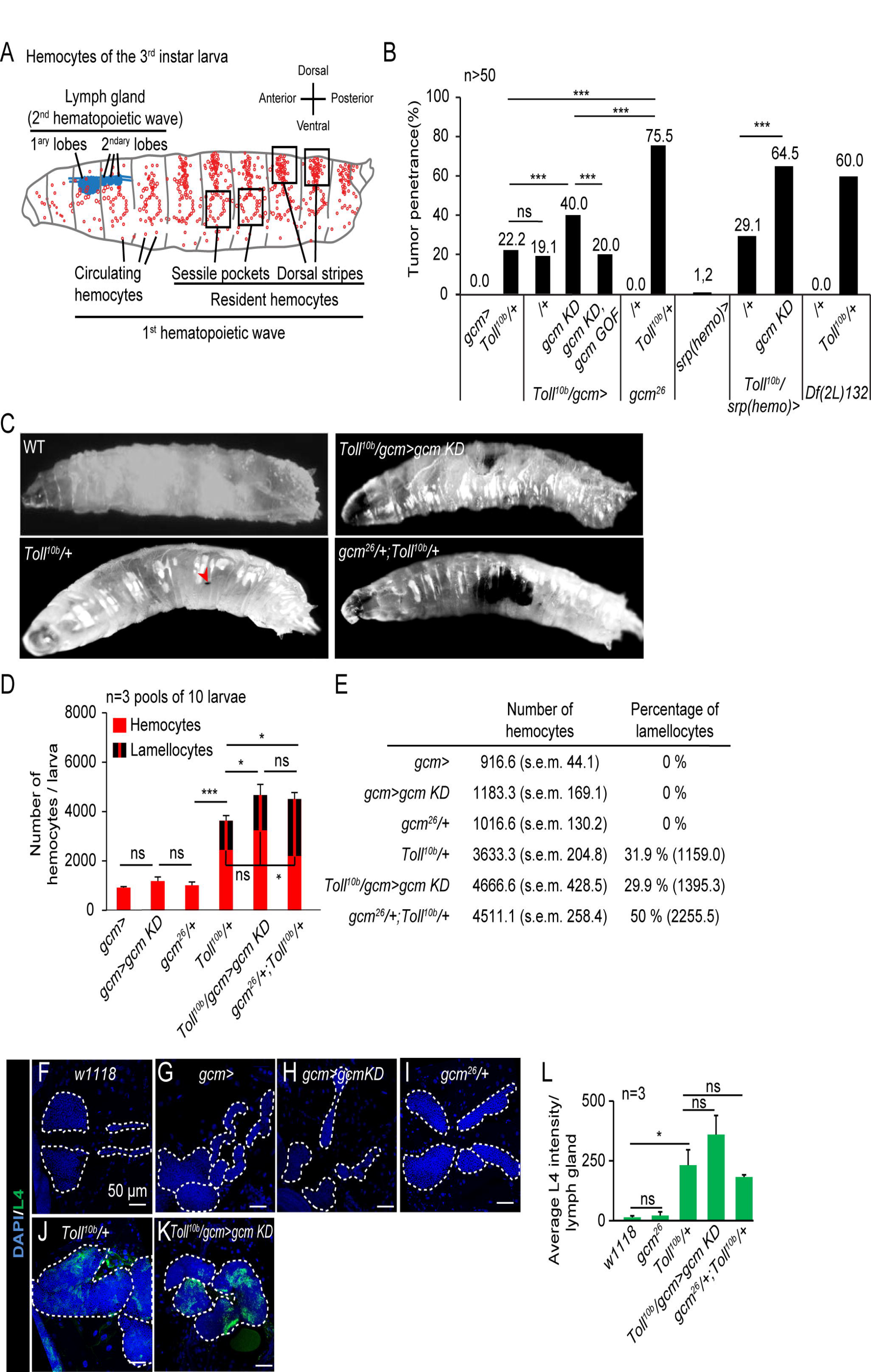
Sensitized hemocytes enhances the inflammatory response induced by Toll activation. **A)** Schematic representation of the hemocytes of the *Drosophila* WL3. The orientation of the larva is indicated on the right. The larva is mostly populated by hemocytes originating from the 1^st^ hematopoietic wave during embryogenesis. In the larva, the embryonic hemocytes (in red) are circulating in the hemolymph or become resident and aggregate between the muscles and the cuticle, laterally around the oenocytes to form the sessile pockets or dorsally to form the dorsal stripes. A second hematopoietic wave is taking place at larval stage in the lymph gland (in blue), composed of successive lobes arranged along the cardiac tube, which produce hemocytes that are release shortly after puparium formation during metamorphosis (77). **B)** Penetrance of melanotic tumors (n>50) in WL3 of the indicated genotype. The p-values were estimated with a chi-square test for frequency comparison. **C)** WL3 of the indicated genotypes. The red arrowhead indicates a small melanotic tumors. **D**,**E)** Total number of hemocytes and lamellocyte contribution (n=3, using 10 larvae/replicate). The p-values were estimated by ANOVA followed by student post hoc test. **F-K)** Lymph glands from WL3 of the following genotype: w1118 (**F**), *gcmGal4 (gcm>*, **G**), *gcmGal4;UAS-gcmRNAi/+* (*gcm>gcm KD*, **H**), *gcm*^*26*^*/+* (**I**), *Toll*^*10b*^*/+* (**J**) and *gcmGal4/+;UAS-gcmRNAi/Toll*^*10b*^ (*Toll*^*10b*^*/gcm>gcm KD*, **K**). The lamellocytes are labelled with an antibody anti-L4 (in green) and the nuclei labelled with DAPI (in blue). The scale bars represent 50 μm. The white dashed lines highlight the lobes of the lymph glands. **L)** Quantification of the L4 intensity in the lymph glands of the indicated genotypes. The p-values were estimated with student test. In all figures, *p<0.0.5, **p<0.01; ***p<0.001, ns: not significant.

The transcription factor Glial cell deficient/Glial cell missing (Glide/Gcm, Gcm throughout the manuscript) is specifically expressed in the hemocytes of the 1^st^ wave and has an anti-inflammatory role that is conserved in evolution (12, 13). Gcm inhibits the pathway by activating Jak/Stat inhibitors (12), raising the question of whether this transcription factor has a general role in the inflammatory response. We here demonstrate that Gcm impacts the Toll pathway. Animals displaying constitutive Toll pathway activation and sensitized hemocytes due to Gcm downregulation produce more lamellocytes than control hemocytes. Transcriptomic analyses reveal that such hemocytes express lower levels of glutathione-S transferase and produce higher levels of reactive oxygen species, which may explain their higher propension to produce lamellocytes. In addition, our data highlight the expression of several neurotransmitter receptors in these sensitized hemocytes, and we show that these receptors regulate the number of hemocytes in the larva.

In sum, the present work indicates that Gcm acts as a general anti-inflammatory transcription factor inhibiting the Toll proinflammatory pathway. Moreover, it highlights a new signalling channel through which neurotransmitters from the nervous system modulate the immune system during inflammation.

## Materials and methods

### Fly strains and genetics

Flies were raised on standard media at 25°C. The stocks used are detailed in supplementary methods

### Monitoring the tumors and hemocyte phenotypes

The tumor, hemocyte and lymph gland phenotypes were scored as in (12). Detailed protocols are available in supplementary methods for the estimation of the penetrance of the melanotic tumours, the hemocytes counting and the lymph gland immunolabelling.

### Stranded RNA sequencing on hemocytes from WL3 larvae

The sample preparation and analysis is detailed in the supplementary methods. The RNA-seq data have been deposited in the ArrayExpress database at EMBL-EBI (www.ebi.ac.uk/arrayexpress) under accession number ***E-MTAB-11970***.

### RNA extraction and qPCR

For the qPCR validation of the transcriptomic data, 20 WL3 of the indicated genotypes were bled on ice cold PBS. The cells were then centrifuged at 1200 rpm at 4°C and RNA isolation was performed with Tri-reagent (Sigma) following the manufacture’s protocol. The DNase treatment was done with the TURBO DNA-free kit (Invitrogen) and the reverse transcription (RT) with the Super-Script IV (Invitrogen) using random primers. The qPCR assays were done with FastStart Essential DNA Green Master (Roche) with the primers are listed in supplementary methods.

The expression levels were calculated relative to the two housekeeping genes Rp49 and Act5C levels using the ΔCt formula: 2^(average(Ct_*Rp49*_, Ct_*Act5c*_) – Ct_target_). Triplicate were done for each genotype and the levels were compared using bilateral student test after variance analysis.

### DHE, pH3 and Dcp-1 quantification

ROS levels were estimated using dihydroethidium (DHE, Sigma) (14). The DHE intensity averages were compared using bilateral student test. For the estimation of the number of mitotic and apoptotic hemocytes, the hemocytes were labelled for pH3 or Dcp-1, respectively. Detailed protocols are available in supplementary methods.

## Results

### Sensitizing hemocytes enhances their response to Toll activation

To understand whether Gcm counteracts the Toll proinflammatory pathway, we combined a *Toll* mutation (15) and altered Gcm expression. *Toll*^*10b*^ is a dominant mutation replacing the amino acid C781Y in the extracellular domain of the receptor, which induces constitutive activation of the Toll pathway (15). Compared to wild type, *Toll*^*10b*^*/+* animals display higher number of hemocytes, precocious lymph glands hystolysis and spontaneous differentiation lamellocytes that aggregate and form melanized black masses called melanotic tumors in 22% of the larvae (**Figure 1B-E**,**F**,**J**)(11, 16-19). In homeostatic conditions, the 3^rd^ larval instar lymph gland is composed of large primary lobes containing progenitors and differentiated hemocytes followed by small secondary lobes composed by undifferentiated hemocytes. In *Toll*^*10b*^*/+* animals, the primary lobes and some secondary lobes are histolyzed, the remaining lobes are enlarged and display mature plasmatocytes and lamellocytes (**Figure 1F**,**J, Supplementary Figure S1A**,**B**) (11).

Knocking down Gcm expression in hemocytes using the gcm-Gal4 driver (*gcm>gcm KD*), does not per se affect hemocyte number or nature (**Figure 1D**,**E**), but *Toll*^*10b*^*/gcm>gcm KD* animals display a considerably enhanced inflammatory phenotype compared to *Toll*^*10b*^*/+* animals. The number of larvae carrying tumors (∼40%) (**Figure 1B**) as well as the number of circulating hemocytes and lamellocytes per larva are significantly higher (**Figure 1D**,**E**). We did not observe significant differences of lamellocytes’ differentiation in the remaining lobes of the double mutant *Toll*^*10b*^*/gcm>gcm KD* compared to *Toll*^*10b*^*/+* lymph glands (**Figure 1F-L**). A similar strengthening of the melanotic tumor phenotype is observed by driving *gcm KD* with the driver *srp(hemo)Gal4 (srp(hemo)> (20)*, **Figure 1B**), which is also specific for the 1^st^ wave hemocytes (21). Importantly, the phenotype of the double mutant animals is rescued by the over-expression of Gcm (*Toll*^*10b*^*/gcm>gcm KD, gcm GOF*) (**Figure 1B**).

The response to Toll activation further increases in combination with the gcm null alleles (*gcm*^*26*^ (22) or the *Df(2L)132* (23)) in heterozygous conditions. *gcm*^*26*^*/+;Toll*^*10b*^*/+* and *Df(2L)132/+;Toll*^*10b*^*/+* display higher penetrance of the melanotic tumor phenotype compared to *Toll*^*10b*^*/+* (**Figure 1B**). As in the case of *gcm KD*, the number of hemocytes in g*cm*^*26*^*/+* animals is not affected, while it does decrease in homozygous embryos (24, 25) (**Figure 1D**,**E**). *gcm*^*26*^*/+, Toll*^*10b*^*/+* animals display similar number of hemocytes but higher proportion of lamellocytes in the hemolymph compared to *Toll*^*10b*^*/gcm>gcm KD*, suggesting an even stronger pro-inflammatory phenotype than *gcm>gcm KD* (**Figure 1B-E**).

In sum, reducing *gcm* expression sensitizes the hemocytes and enhances the response to Toll pathway activation.

### Transcriptome analysis of the sensitized hemocytes after Toll pathway activation

To assess the molecular mechanisms underlying the relative impact of Toll and Gcm on the observed phenotypes, we performed pairwise comparisons amongst the transcriptomes from *gcm*^*26*^*/+*, from *Toll*^*10b*^*/+* and from *gcm*^*26*^*/+;Toll*^*10b*^*/+* wandering 3^rd^ instar larvae (WL3) hemocytes (**Supplementary Figure S2A**,**B**).

The comparison of *gcm*^*26*^*/+;Toll*^*10b*^*/+* with *gcm*^*26*^*/+* hemocytes highlights the impact of *Toll*^*10b*^ on gene expression: 688 genes are significantly upregulated (mean expression > 100, Log2FC > 1 and p < 0.01) (**Figure 2A, Supplementary Table S1**). In line with the known function of the Toll pathway in response to fungi, bacteria and wasp infestation (26-28), Gene Ontology analysis indicates the upregulation of genes involved in the innate immune response and more specifically in the Jak/Stat pathway, in the defense response to Gram-positive and Gram-negative bacteria (**Supplementary Figure S2C**). The expression of most core components of the Toll pathway is induced, including that of the transcription factor *dorsal (dl)* (**Figure 2A’, Supplementary Figure S2F**), in agreement with the autoregulatory loop shown for the Toll pathway (29). The induction of the core elements of the Jak/Stat pathway (**Supplementary Figure S2D**) is concordant with ChIP data targeting Dorsal (Dl), which indicates that Dl binds the promoters of all Jak/Stat core components (30). The response to Gram-negative bacteria is commonly associated with the activation of the IMD pathway and illustrates the cross-talk between the Toll and the IMD pathways (31-34). Most core components of the IMD pathways as well as the majority of anti-microbial peptides are upregulated in *gcm*^*26*^*/+;Toll*^*10b*^*/+* compared to *gcm*^*26*^*/+* (**Figure 2A’-A’’’**,**B**), suggesting that the Toll pathway may activate the IMD pathway. Concordantly, ChIP data targeting Dl and Dorsal-related immunity factor (Dif) show that most genes of the IMD pathway are targeted by Dl/Dif in the larva (35) and a transcriptome analysis of *Toll*^*10b*^ animals shows that Relish (Rel) is induced in *Toll*^*10b*^ adults (32).

**Figure 2:**
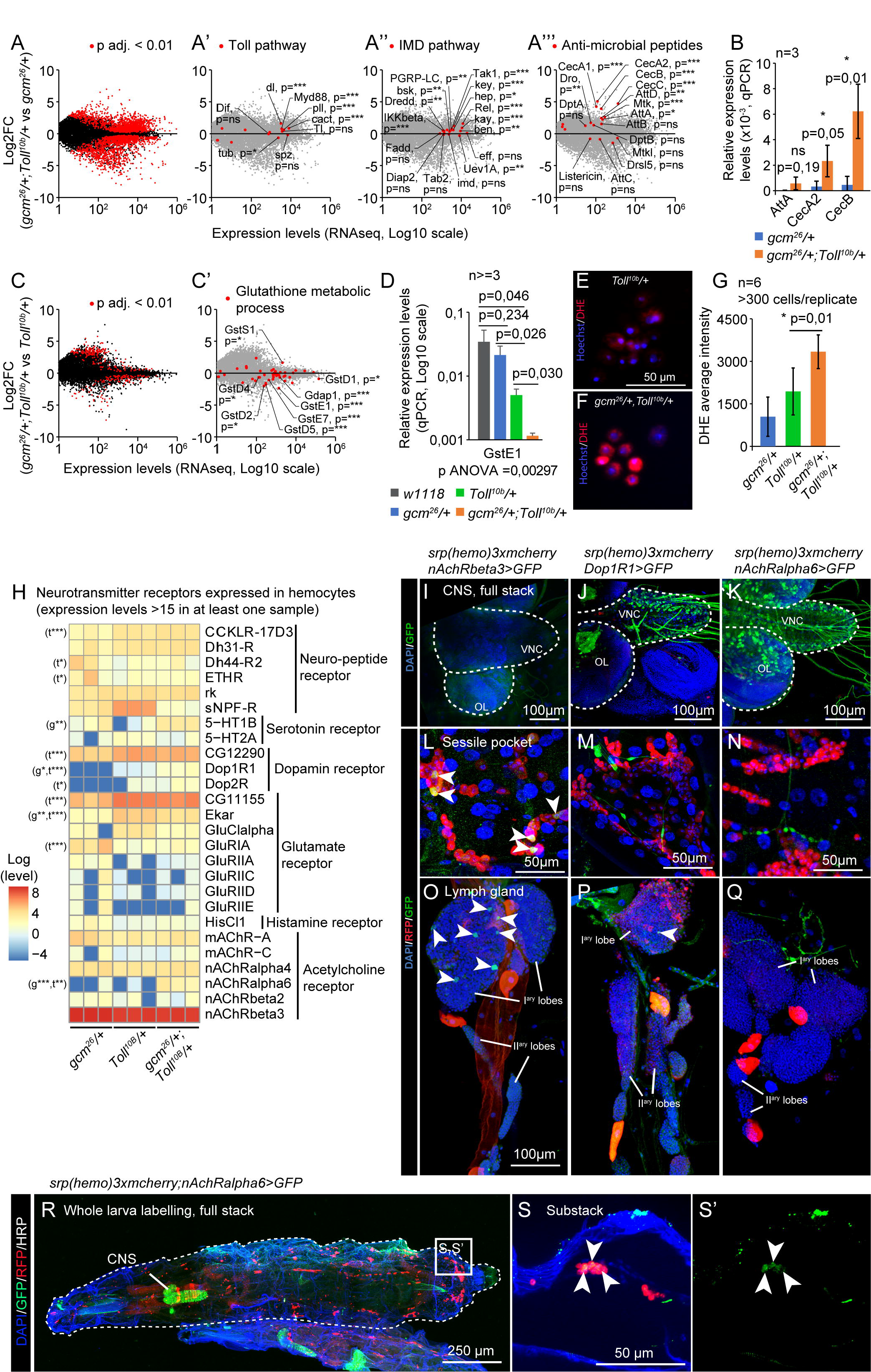
The proinflammatory condition *gcm*^*26*^*/+;Toll*^*10b*^*/+* induces the IMD pathway, modulate the ROS metabolism and the expression of neurotransmitter receptors. **A-A’’’)** Transcriptome comparison of hemocytes from WL3 *gcm*^*26*^*/+* and *gcm*^*26*^*/+;Toll*^*10b*^*/+*. The x-axis represents the average genes expression levels (n=3) and the y-axis the Log2 Fold Change *gcm*^*26*^*/+;Toll*^*10b*^*/+ / gcm*^*26*^*/+*. The red dots highlight the genes presenting significant fold change (adjusted p-values < 0.01) in **(A)**, the genes of the Toll pathway in **(A’)**, of the IMD pathway in **(A’’)** and the genes coding for antimicrobial peptides (AMP) in **(A’’’)**. **B)** Expression levels of *AttA, CecA2* and *CecB* in hemocytes *gcm*^*26*^ (in blue) and *gcm*^*26*^*/+;Toll*^*10b*^*/+* (in orange) estimated by quantitative PCR. N = 3 pools of 10 larvae, p-value estimated by bilateral student test. **C**,**C’)** Transcriptome comparison of hemocytes from WL3 *Toll*^*10b*^ and *gcm*^*26*^*/+;Toll*^*10b*^*/+*. The x-axis represents the average genes expression levels (n=3) and the y-axis the Log2 Fold Change *gcm*^*26*^*/+;Toll*^*10b*^*/+/Toll*^*10b*^. The red dots highlight the genes presenting significant fold change (adjusted p-values < 0.01) in **(C)** and the genes coding for glutathione-S-transferases (Gst) in **(C’)**. **D)** Expression levels of *GstE1* in hemocytes *w1118* (wild type control, in gray), *gcm*^*26*^*/+ (in blue), Toll*^*10b*^*/+* (in green) and *gcm*^*26*^*/+;Toll*^*10b*^*/+* (in orange) estimated by quantitative PCR. N >= 3 pools of 10 larvae, p-value estimated by bilateral student test after ANOVA (p ANOVA = 0,00297). **E-G)** Live hemocytes from *Toll*^*10b*^*/+* **(E)** and *gcm*^*26*^*/+;Toll*^*10b*^*/+* **(F)** animals labelled for reactive oxygen species (ROS) using DHE (in red). The nuclei are labelled with Hoechst. The levels of oxidised DHE are quantified in **(G)**. N = 6, at least 300 hemocytes were monitored per replicate, the p-value was estimated by bilateral student test. **H)** Heatmap representing the expression levels of neurotransmitter and neuropeptide receptors in hemocytes *gcm*^*26*^*/+, Toll*^*10b*^*/+* and *gcm*^*26*^*/+;Toll*^*10b*^*/+*. The levels are in log scale. The significant p-values are mentioned on the left side of the heatmap. The comparison of *gcm*^*26*^*/+* to *gcm*^*26*^*/+;Toll*^*10b*^*/+* is indicated by “t” and *Toll*^*10b*^*/+* to *gcm*^*26*^*/+;Toll*^*10b*^*/+* “g”, p-values: *:0.05<p<0.01; **: 0.01<p<0.001; ***: 0.001<p. **I-Q)** Central nervous systems (**I-K**), sessile pockets (**L-N**) and lymph glands (**O-Q**) from larvae carrying the *T2A-Gal4* reporters of *nAchRbeta3, Dop1R1* or *nAchRalpha6* (in green) and the hemocyte reporter *srp(hemo)3xmcherry* (in red). The nuclei are labelled with DAPI. The images were acquired with confocal microscopy and represent the whole stack projections. Scale bars are 100 μm in **(I-K**,**O-Q)** and 50 μm in **(L-N)**. **R**,**S)** Whole mount immunolabellings of L3 larvae *nAchRalpha6-T2A-Gal4/srp(hemo)3xmcherry;UAS-encGFP/+*. The larva is outlined by a white dashed line, the hemocytes are labelled with anti-RFP (in red) and the cells expressing *nAchRalpha6-T2A-Gal4* with anti-GFP (in green). The complete stack projections are shown in **(R)**, a substack of the region indicated in **(R)** is shown in **(S**,**S’)**.

The impact of hemocyte sensitization is shown by comparing the transcriptomes from *gcm*^*26*^*/+;Toll*^*10b*^*/+* and *Toll*^*10b*^*/+* larvae: 87 genes are down-regulated and 161 genes are upregulated by *gcm*^*26*^ (mean expression > 100, absolute value [Log2FC] > 1 and p < 0.01) (**Supplementary Table S1**). Noteworthily, the number of genes affected by *gcm*^*26*^ is much lower than that affected by *Toll*^*10b*^ (**Figure 2A**,**C**). This is likely due to the fact that *Toll*^*10b*^ is a dominant, gain of function condition while *gcm*^*26*^ is a recessive mutation analyzed in heterozygous conditions, thus, a stronger impact on the transcriptome is expected for *Toll*^*10b*^. With such low number of genes, only few GO term were found significantly enriched when *gcm*^*26*^*/+;Toll*^*10b*^*/+* and *Toll*^*10b*^*/+* larvae were compared. We did follow one of the GO terms with the lowest p-value, glutathione metabolic process (**Supplementary Figure S2E**), and found that most associated genes are down-regulated in *gcm*^*26*^*/+;Toll*^*10b*^*/+* compared to *Toll*^*10b*^*/+* hemocytes (**Figure 2C’**,**D**) and belong to the glutathione S transferase family (Gst). Since Gsts are involved in xenobiotic detoxification and in the defense mechanism against oxidative stress (36-38), we assessed the biological relevance of their reduction in hemocytes, by estimating the ROS levels in *gcm*^*26*^*/+;Toll*^*10b*^*/+* and *Toll*^*10b*^*/+* hemocytes using dihydroethidium. Dihydroethidium is oxidized by intracellular ROS to produce red fluorescent ethidium (39). The quantification of the ethidium levels in hemocytes suggests higher levels of ROS in *gcm*^*26*^*/+;Toll*^*10b*^*/+* than in Toll^10b^/+ hemocytes, which can be due to the lower levels of Gst (**Figure 2E-G**). A recent study showed that the ROS produced after injury induces the Toll pathway in hemocytes (40). Thus, we speculate that the low levels of Gst in *gcm*^*26*^*/+;Toll*^*10b*^*/+* hemocytes (**Figure 2D**) may increase their level of ROS, hence leading to a stronger response to Toll activation.

Unexpectedly, the GO term enrichment analyses carried out on each dataset highlighted GO term related to synapse activity enriched in the two comparisons (**Supplementary Figure S2C**,**E**). These genes included several neurotransmitter receptors, suggesting an involvement of neurotransmitter mediated signaling in Toll activation and *gcm*^*26*^ sensitization. A targeted analysis of neurotransmitter receptor expression in our dataset revealed the expression of 26 receptors in hemocytes and significant upregulations of the Nicotinic Acetylcholine receptor alpha 6 (nAchRalpha6), the serotonin receptor 5-hydroxytryptamine receptor 1B (5-HT1B), the dopamine receptor Dopamine 1-like receptor 1 (Dop1R1) and the short neuropeptide F receptor (sNPF-R) by *Toll*^*10b*^ and/or by *gcm*^*26*^ (Figure 2H, Supplementary Table S1). nAchRalpha6, 5-HT1B and Dop1R1 expression is significantly enhanced in the double mutant *gcm*^*26*^*/+;Toll*^*10b*^*/+* compared to *Toll*^*10b*^*/+* and to *gcm*^*26*^*/+*, with nAchRalpha6 showing the strongest increase. In contrast, sNPF-R is significantly reduced (**Figure 2H, Supplementary Table S1**). Other neurotransmitter receptors such as nAchRbeta3 are expressed constitutively at high levels in the hemocytes regardless of the genetic background (**Figure 2H**).

To verify the expression of neurotransmitter receptors in hemocytes, we took advantage of recently produced T2A reporter lines that express intact receptors along with Gal4 under the endogenous promoter of the gene (41, 42). We assessed the expression of *nAchRbeta3*, which presents the highest expression levels and is constitutively expressed in hemocytes (**Figure 2H**), as well as *Dop1R1* and *nAchRalpha6*, which show the most significative induction in the double mutant *gcm*^*26*^*/+;Toll*^*10b*^*/+* compared to *Toll*^*10b*^*/+* and *gcm*^*26*^*/+*. The *T2A-Gal4* lines were crossed with the double reporter *srp(hemo)-3xmcherry;UAS-GFP* to obtain flies that expresses RFP in hemocytes (both lymph gland and 1^st^ wave hemocytes) (43) and GFP in the receptor-*T2A-Gal4* expressing cells. The *Dop1R1* and *nAchRalpha6* reporters but not *nAchRbeta3* are expressed in the larval CNS (**Figure 2I-K**), consistent with the literature (41, 44).

The *nAchRbeta3* reporter shows GFP signals in hemocytes from the lymph gland and in sessile pockets (**Figure 2L**,**O**) and the *Dop1R1* reporter is detected in few cells of the lymph gland but not in the hemocytes of the sessile pockets (**Figure 2M**,**P**), nor in other circulating or resident hemocytes. The *nAchRalpha6-T2A-Gal4* reporter is not detected in the sessile pockets nor in the lymph gland (**Figure 2N**,**Q**), however, whole larva immunolabelling and larval fillet preparations show expression of the receptor in resident hemocytes located in the dorsal stripes (**Figure 1A, Figure 2R**,**S, Supplementary Figure S3A-C**). The larva contains on average 1.05% +/- 0.49 of nAchRalpha6 positive hemocytes (n=3, estimated by cytometry on 10 larvae per replicate). Because of the highest effect observed in the double mutant larvae, we focused on *nAchRalpha6* and confirmed its expression profile with a transgenic line expressing a nAchRalpha6-YFP fusion protein (45) (**Supplementary Figure S3D**,**E**). The nAchRalpha6 positive hemocytes express strongly the plasmatocyte markers Nimrod C1 (NimC1 or P1) and Hemese (He) (**Supplementary Figure S3C-E’’’**).

Overall, these data suggest that Toll activation regulates the IMD and the Jak/Stat pathway, that sensitized hemocytes display higher ROS levels in response to Toll activation possibly due to suboptimal levels of Gst, and that the hemocytes express neurotransmitter receptors, whose expression is modulated by inflammatory conditions.

### nAchRalpha6 modulates the proliferation of hemocytes

We next evaluated the impact of the receptors on hemocytes. We focused on *nAchRalpha6* and observed how manipulating its expression levels affects hemocytes. The null mutation *nAchRalpha6*^*DAS1*^ alters the splice donor site of the first intron, which produces an inactive truncated protein, the null mutation *nAchRalpha6*^*DAS2*^ converts the codon for the tryptophan 458 to a terminal codon. In both mutations, the number of hemocytes in WL3 is significantly reduced (**Figure 3A**). Given the impact of nAchRalpha6 in the nervous system (46), we next determined if the phenotype is cell autonomous by downregulating the expression of nAchRalpha6 specifically in hemocytes. The expression of a UAS-RNAi transgene targeting the receptor was driven by a combination of the two larval hemocyte specific drivers *HmldeltaGal4* and *PxnGal4* that cover the whole larval hemocyte population (47-49). The *nAchRalpha6* knock down (*nAchRalpha6-KD*) animals are completely viable and display fewer hemocytes than the control animals (**Figure 3B**). The hemocyte number is also reduced in *nAchRalpha6-KD* with the driver *PxnGal4* alone (**Figure 3C**), but not with *HmldeltaGal4* alone (**Figure 3D**). The two drivers are specific to hemocytes and while the majority of hemocytes express both drivers, small subsets of hemocytes express exclusively *PxnGal4* (37% in WL3) or *HmldeltaGal4* (10% in WL3) (47). The different hemocyte number observed upon *nAchRalpha6 KD* driven by one or the other driver may depend either on the different hemocyte populations affected or on the different levels of knock down. To discern between the two possibilities, we stabilized and hence enhanced *HmldeltaGal4* driven expression levels using the G-trace approach (50). *HmldeltaGal4,Dbgtrace>nAchRalpha6-KD* animals do display fewer hemocytes compared to the control WL3 (**Figure 3E**), indicating that the difference observed upon *PxnGal4* and *HmldeltaGal4* driven knock down is due to different levels of Gal4 induction.

**Figure 3:**
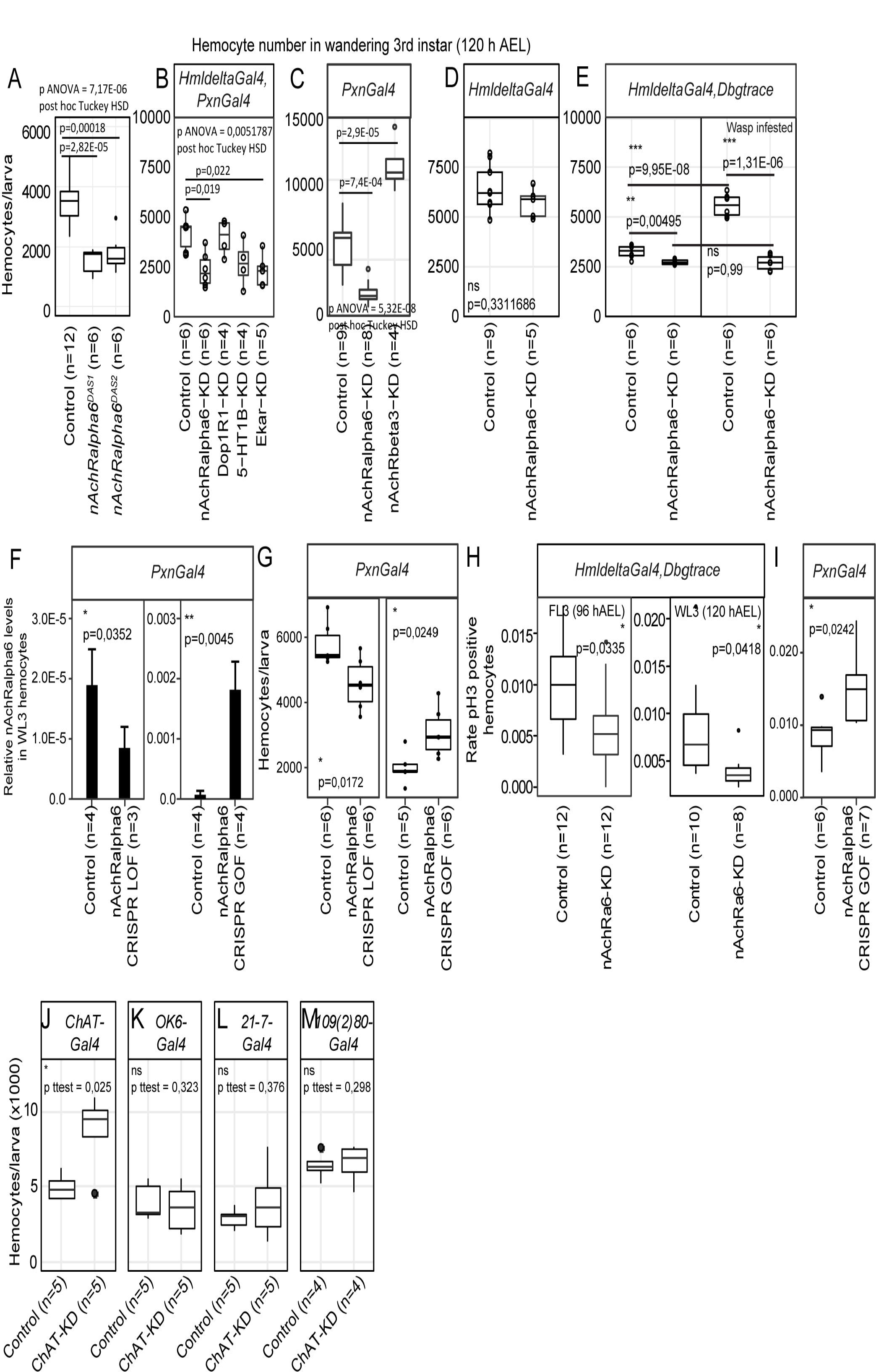
nAchRalpha6 modulates hemocytes proliferation, cell autonomously. **A-E)** Number of hemocytes per WL3 in the indicated genotypes. N => 4, each replicate being a pool of 10 females. Note that in *HmldeltaGal4,Dbgtrace* (**E**), Gal4 expression was enhanced using lineage tracing *Gal4* transgenes (*Dbgtrace*). *Dbgtrace* includes a flipase cassette under the control of the UAS promoter (UAS-FLP) and the Gal4 gene separated from the Act5C constitutive promoter by a stop cassette surrounded by two flipase recognition sites (*Act5C-FRT-STOP-FRT-Gal4*) (50). The expression of the flipase in the hemocytes excises the STOP cassette and leads to constitutive expression of *Gal4* in those cells, strongly enhancing the expression levels of the Gal4. **F)** Expression levels of *nAchRalpha6* in WL3 hemocytes from *Pxn>nAchRalpha6 CRISPR LOF* and *GOF* animals (complete genotypes indicated in the method section) measured by quantitative PCR; n = 4 each replicate being a pool of 10 females. Note that the controls are specific to each genetic setup. The full genotypes of the controls are indicated in the method section “Fly strains and genetics”. In *CRISPR LOF* animals, the Cas9 nuclease was expressed specifically in hemocytes using the driver *PxnGal4* and targeted to the coding sequence of nAchRalpha6 by the constitutive expression of two *nAchRalpha6* specific guide RNA (*nAchRalpha6 CRISPR LOF*) (78). In *CRISPR GOF* animals, a dead Cas9 fused to the activator domain VPR (Z_IRIN_ et al. 2020) was expressed with the driver *PxnGal4* and guided to the promoter of nAchRalpha6 with specific guide RNA (nAchRalpha6 CRISPR GOF). **G)** Number of hemocytes per WL3 in *Pxn>nAchRalpha6 CRISPR LOF* and *GOF*. N => 5, each replicate being a pool of 10 females. **H)** Quantification of the proliferative hemocytes in feeding L3 (96 hAEL) and WL3 (120 hAEL) *HmldeltaGal4,Dbgtrace* or *HmldeltaGal4,Dbgtrace,nAchRalpha6-KD*; n =>8, with more than 300 hemocytes scored for each replicate, p-values were estimated by one-factor ANOVA. **I)** Quantification of the proliferative hemocytes in WL3 (120 hAEL) *Pxn>nAchRalpha6 CRISPR GOF* and control; n =>6, with more than 1000 hemocytes scored for each replicate, p-value was estimated by one-factor ANOVA. **J-M)** Number of hemocytes per WL3 (120 hAEL) in the indicated genotypes. ChAT expression was inhibited specifically in cholinergic neurons with ChATGal4 **(J)**, in type I motoneurons with OK6Gal4 **(K)** and in multidendritic neurons with 21-7Gal4 or 109(2)80Gal4 **(L**,**M)**. N => 4, each replicate being a pool of 10 females.

To further prove the role of *nAchRalpha6* in hemocytes, we used tissue specific CRISPR/Cas9 mediated loss of function (LOF) and gain of function (GOF) animals (**Figure 3F**,**G**). In *nAchRalpha6 CRISPR LOF* larvae, the expression levels of nAchRalpha6 decrease and so does the number of hemocytes (**Figure 3F**,**G**). In nAchRalpha6 CRISPR GOF, the expression levels of nAchRalpha6 increase by more than 10 folds and the number of hemocytes increase compared to the control (**Figure 3F**,**G**). Altogether, these data demonstrate that nAchRalpha6 regulates the number of hemocytes in the larva.

To assess the cause(s) of the different hemocyte number in *nAchRalpha6 KD* and *GOF*, we quantified hemocyte proliferation and apoptosis using antibodies against phosphorylated Ser10 of histone 3 (pH3) (51) and cleaved caspase Dcp-1 (52), respectively. No difference was found in the rate of apoptosis (**Supplementary Figure S4B-D**). A significant reduction of the proliferation rate was observed in *nAchRalpha6-KD* hemocytes in feeding as well as in wandering 3^rd^ instar larvae (FL3, WL3 respectively) and a significant increase in proliferation in *nAchRalpha6 CRISPR GOF* hemocytes, compared to control hemocytes (**Figure 3H**,**I Supplementary Figure S4A**). To assess if the impact of nAchRalpha6 on proliferation is cell autonomous, we quantified the number of proliferative hemocytes that express nAchRalpha6 in *nAchRalpha6-T2A-Gal4/+;UAS-GFP/srp(hemo)3xmcherry* larvae. On average, 26.6% +/- 2.6 of nAchRalpha6 hemocytes are mitotic according to pH3 labelling compared to 0.6% in the whole population (n=3, estimated on 10 larvae per replicate, p paired student test = 0.0038). The nAchRalpha6 hemocytes represent 46% +/- 6.5 of the proliferative larval hemocytes, suggesting that the receptor is involved in the cell autonomous regulation of hemocyte proliferation. These results are in agreement with the increase in hemocyte number observed in the *gcm*^*26*^*/+;Toll*^*10b*^*/+* larvae that also display increased expression of *nAchRalpha6*. We hence assessed the direct impact of nAchRalpha6 on hemocyte number in inflammatory conditions. Given the complexity of the genetic set up to reduce nAchRalpha6 expression in *Toll*^*10b*^ animals, we induced an immune challenge upon infesting larvae with the parasitoid wasp *Leptopilina boulardi*, which is known to trigger the Toll pathway. The rate of lamellocyte differentiation is not affected in *HmldeltaGal4,Dbgtrace>nAchRalpha6-KD* animals, with a percentage of lamellocytes/total hemocytes number of 18.6% (n = 6, stdev. = 2.9%), compared to 18.4% in control animals (n = 6, stdev = 12.3%; p t test unequal variance = 0.97). However, a striking difference is observed in the number of hemocytes. Control animals display significantly more hemocytes after wasp infestation, while the hemocyte number in *nAchRalpha6-KD* animals remains stable (**Figure 3E**). Thus, the increase of hemocyte triggered by wasp infestation depends at least in part, on *nAchRalpha6* expression in hemocytes.

Overall, these data show that *nAchRalpha6* modulates the proliferation of hemocytes in homeostasis and during the inflammatory response.

### Cholinergic signaling regulate hemocyte homeostasis

We evaluated the impact of other neurotransmitter receptors and down-regulated 5-HT1B, Dop1R1, nAchRbeta3 or the glutamate receptor Eye-enriched kainate receptor (Ekar) (**Figure 3B**). In all cases, the KD animals are completely viable (data not shown). In terms of hemocyte number, *5-HT1B-KD* and *Dop1R1-KD* show no difference compared to control WL3. *Ekar-KD* displays less hemocytes (**Figure 3B**), suggesting that glutamate signaling may also be involved in the regulation of hemocyte homeostasis. *nAchRbeta3-KD* shows a strong increase in the number of hemocytes (**Figure 3C**), which further highlights the importance of cholinergic signaling in hemocytes’ homeostasis.

At last, to assess the impact of cholinergic transmission on hemocytes, we monitored the number of larval hemocytes after inhibiting the expression of the choline-acetyltransferase ChAT in neurons. To inhibit ChAT, we used a *UAS-ChAT-RNAi* (53) and a *ChAT-Gal4* to drive *ChAT-RNAi* in the cholinergic neurons (54), which led to a strong increase in the number of hemocytes (**Figure 3J**), indicating that cholinergic neurons modulate hemocyte homeostasis through secretion of acetylcholine. As controls, we used *OK6-Gal4* for type 1 motoneurons (55), which mostly secrete glutamate (56, 57), *21-7-Gal4* and 109(2)*80-Gal4* drivers for multidendritic neurons, which are cholinergic (54) and regulate hemocyte localisation and proliferation at the dorsal stripes through activin signaling (58). Since the multidendritic neurons make synapses at the central nervous system (CNS) (59-62), we do not expect an effect using these drivers either (**Figure 3K-M**). These data indicate that motoneurons or multidendritic neurons are not the source of acetylcholine that affects hemocyte number.

Taken together, our data show that several neurotransmitter receptors are involved in hemocyte homeostasis, that nAchRalpha6 regulates the proliferation of the hemocytes cell autonomously and that acetylcholine signaling to the hemocytes likely originates from cholinergic neurons from the CNS.

## Discussion

In this study, we show that downregulating Gcm enhances the immune response to Toll activation, calling for a general anti-inflammatory role of this evolutionarily conserved transcriptional cascade. The comparison of the transcriptomes in control and mutant backgrounds reveals that the activation of the Toll pathway induces the expression of core components of the IMD pathway and that sensitizing the hemocytes by Gcm downregulation alters the levels of Gst and ROS metabolism in *Toll*^*10b*^ background. Finally, we demonstrate that hemocyte expression of acetylcholine receptor nAchRalpha6 is modulated upon hemocyte sensitizing and Toll activation and that nAchRalpha6 regulates hemocyte proliferation cell-autonomously. The finding that cholinergic signaling controls hemocyte proliferation underlines the interaction between immune and nervous systems.

### Sensitized hemocytes display an enhanced response to Toll and Jak/Stat signaling

Gcm acts as a general anti-inflammatory factor as its downregulation enhances the inflammatory response to challenges of different nature. This phenotype is observed in sensitized hemocytes upon the constitutive activation of the Toll pathway (this study) or of the Jak/Stat pathway (Bazzi et al., 2018), two examples of chronic challenge. A similar phenotype is also observed upon wasp infestation, an acute challenge that activates both pathways (12, 28). The inflammatory responses induced by Toll and Jak/Stat are highly similar: increasing hemocyte number and lamellocyte differentiation at comparable levels (12). In both conditions, sensitizing the hemocytes doubles the penetrance of the melanotic tumor phenotype in the larva (12). These strong similarities can be explained by the high interconnection between the two pathways. For example, the Toll pathway acts upstream of Jak/Stat for the regulation of the thiolester-containing protein Tep1 (63). In addition, the Tep protein family regulates the Toll pathway (64) and Jak/Stat modulates the expression of the Toll’s ligand Spatzle (65), providing means by which the Jak/Stat pathway can modulate the Toll pathway. At last, recent data show that in hemocytes, Toll activation induces the expression of the pro-inflammatory cytokine Upd3, which activates the Jak/Stat pathway (66). Therefore, the activation of one pathway will likely activate the second one in a feed-forward loop. This hypothesis is supported by the *Toll*^*10b*^ transcriptome that shows increased levels of several targets regulated by the Jak/Stat pathway, including Ptp61F and Socs36E (67)(**Supplementary Figure S2D**) as well as Tep1, Tep2 and Tep4 (63, 65)(**Supplementary Table 1**).Our previous data showed that Gcm inhibits the Jak/Stat pathway (12). Gcm could hence inhibit the Toll pathway at least in part through the inhibition of the Jak/Stat pathway.

A second hypothesis that can explain the impact of Gcm on the inflammatory response is the modulation of the Gst. Our transcriptome analysis on the sensitized animals reveals a significant decrease in the anti-oxidant enzymes Gst, which correlates with higher levels of ROS. The production of ROS is tightly linked to the Toll pathway. On the one hand, ROS are known to activate the Toll and the Jak/Stat pathways (40, 68), on the other hand, Toll activates the production of ROS (69). We speculate that in our sensitized model, the deficit in Gst increases ROS levels, which might enhance the inflammatory response induced by Toll activation.

This data highlight Gcm as a potent anti-inflammatory transcription factor acting at multiple levels, directly on the Jak/Stat pathway, indirectly on Toll and IMD pathway or on ROS levels through the modulation of the Gst. Importantly, the impact of Gcm on the inflammatory response of immune cells is conserved in mammals. In mouse, knock out for Gcm2 in microglia, the macrophage of the nervous system, leads to the production of microglia in a pro-inflammatory state (13).

### Neurotransmitter receptors’ expression in hemocytes

The modulation of the immune cells by neurotransmitters is well described in mammals. Numerous neurotransmitter receptors are expressed in immune cells and cholinergic, dopaminergic and serotoninergic signaling mediate the function, the inflammatory status and the proliferation of macrophages (reviewed by (70)). In *Drosophila*, few studies report neurotransmitter signaling in immune cells. Neuronal GABA is secreted in the hemolymph after olfactory stimuli induced by parasitoid wasp scent and promote the differentiation of lamellocytes in the lymph gland (71). Qi et al. (72) showed the impact of serotonin signaling on the phagocytic capacity of the plasmatocytes in the butterfly Pieris rapae and in *Drosophila*. Immune challenge in the adult induces the secretion of serotonin by the plasmatocytes, which enhances their phagocytic capacity. This autocrine process is mediated by the receptors 5-HT1B and 5-HT2B (72). At last, Dopamine signaling is used by the progenitors in the lymph gland to regulate cell cycle (73).

Our transcriptome analysis reveals the expression of a dozen neurotransmitter receptors in the hemocytes, some of whose appear to be modulated by the inflammatory state of the larva. We report here the expression of receptors to acetylcholine, glutamate, serotonin, dopamine and several neuropeptides in the hemocytes patrolling the larva. Our data indicate that the levels of nAchRalpha6 increase in sensitized hemocytes in *Toll*^*10b*^ background and that nAchRalpha6 is enriched in proliferative hemocytes. Additionally, we have shown that modulating acetylcholine production in the nervous system or the expression of specific subunits of the acetylcholine receptors in the hemocytes has a significant impact on these cells. Repressing cholinergic signaling from the neurons increases the number of hemocytes, similarly to the effect of inhibiting *nAchRbeta3* or over-expressing *nAchRalpha6* in hemocytes and opposite to the effect of inhibiting *nAchRalpha6* in hemocytes (**Figure 3C**). Taken together, these data indicate that cholinergic signaling regulate the proliferation of hemocytes through the activation of nicotinic acetylcholine receptors. The nicotinic acetylcholine receptor is composed of five subunits homomeric or heterodimeric (74) and the composition of the receptor defines its biochemical properties (75). Our experimental setup modulates the expression of specific subunits, which may modify the composition of the receptors in the hemocytes and lead to two distinct effects (*i*.*e*. promotion or inhibition of hemocyte proliferation with nAchRalpha6 or nAchRbeta3, respectively). Thus, modulating the expression of neurotransmitter receptor subunits may represent a novel mechanism by which the hemocyte homeostasis is regulated in response to pro-inflammatory cues.

Altogether, these observations suggest that hemocytes are sensitive to a large panel of neurotransmitters. Our data do not allow to distinguish if the signal is transmitted through direct neuron-hemocyte connection or through systemic acetylcholine secretion. Several neurotransmitters are secreted systemically in the hemolymph (71) and can be produced by the hemocytes themselves (72, 73), but this was never shown for acetylcholine in the hemolymph and our transcriptomic data indicate that ChAT is not expressed in hemocytes. A recent study in adult *Drosophila* shows that other tissues than neurons produce acetylcholine and that both neuronal and glia derived acetylcholine regulates the Toll mediated immune response of hemocytes through nAch receptor (76). Thus, cholinergic signaling appears as a fundamental mechanism of the immune response, providing a direct communication channel between the nervous system and the immune system. Establishing the prevalence, the localization and the nature of the receptors expressed in the hemocytes as well as the source of the neurotransmitters will be key steps to decipher this signaling axis.

### Concluding notes

Our study ascertains the anti-inflammatory role of Gcm on several inflammatory pathways, reveals a role for nAchRalpha6 in the regulation of hemocyte proliferation in homeostasis as well as in response to inflammation and shows the contribution of the neuronal cholinergic signaling to the immune system homeostasis. These data parallel the function of neurotransmitter receptors in mammals, whose activation in macrophages modulates cell proliferation and the activity of inflammatory pathways. Our model paves the way to characterize the role of neurotransmitter signaling in the immune response and to explore the evolutionary conserved mechanisms involved.

## Acknowledgments

We thank K. Bruckner, D. Siekhaus, D. Hultmark, H. Tanimoto, S. Kondo, S. Russell, V. Honti, I. Ando and P. Soba for providing fly stocks and antibodies. In addition, stocks obtained from the Bloomington *Drosophila* Stock Center (NIH P40OD018537) and antibodies obtained from the Developmental Studies Hybridoma Bank created by the NICHD of the NIH and maintained at The University of Iowa (Department of Biology, Iowa City, IA 52242) were used in this study. We thank F. Stenger, M. Heinis, M. Moog, A. Kucan, A. Bhan, E. Naegelen, G. Zhang for technical assistance. We thank the Imaging Center of the IGBMC for technical assistance. We thank A. Maglott-Roth from the IGBMC screening facility for the hemocyte analysis. The sequencing was performed by the GenomEast platform, a member of the “France Génomique” consortium (ANR-10-INBS-0009). This work was supported by INSERM, CNRS, UDS, Ligue Régionale contre le Cancer, Hôpital de Strasbourg, ARC, CEFIPRA, USIAS, FRM and ANR grants. P Cattenoz was funded by the ANR and by the ARSEP, W Bazzi by the USIAS and by the FRM (FDT20160435111). S Monticelli was funded from CEFIPRA and ANR fellowships. This work of the Interdisciplinary Thematic Institute IMCBio, as part of the ITI 2021-2028 program of the University of Strasbourg, CNRS and Inserm, was supported by IdEx Unistra (ANR-10-IDEX-0002), and by SFRI-STRAT’US project (ANR 20-SFRI-0012) and EUR IMCBio (ANR-17-EURE-0023) under the framework of the French Investments for the Future Program.

