## Supplementary figures and methods for "Gcm alleviates the inflammatory phenotype induced by Toll activation in *Drosophila* hemocytes"

#### Supplementary Figures, method and table
#### “Gcm alleviates the inflammatory phenotype induced by Toll activation in *Drosophila* hemocytes”

Authors:

Wael Bazzi 1,2,3,4
Sara Monticelli 1,2,3,4
Claude Delaporte 1,2,3,4
Céline Riet 1,2,3,4
Angela Giangrande 1,2,3,4,§
Pierre B. Cattenoz 1,2,3,4,§

List of files :

Supplementary Figure S1: priming hemocytes enhance the melanotic tumors induced by Toll.

Supplementary Figure S2: Transcriptomic analysis of sensitized hemocytes in Toll10b background.

Supplementary Figure S3: nAchRalpha6 is expressed in resident hemocytes.

Supplementary Figure S4: Quantification of apoptotic hemocytes in nAchRalpha6 knock down and gain of function animals.

Supplementary methods

#### Supplementary Figure S1: priming hemocytes enhance the melanotic tumors induced by Toll.

**A,B)** Lymph glands from *w1118* (**A**) and *Toll$^{10b}$/+* (**B**) WL3 labelled with anti-L4 (lamellocyte marker in green), anti-Pxn (mature hemocyte marker in red) and DAPI. The lobes are indicated with white dashed lines.

**C)** Expression levels of the anti-microbial peptides AttB, AttA and CecB, of the cytokines Upd2 and Upd3 as well as of inhibitors and downstream targets of the Jak/Stat pathway in hemocytes *w1118* (in gray) and *gcm$^{26}$/+* (in blue) estimated by quantitative PCR. N = 4 pools of 10 larvae, the ANOVA indicated no significant modulation between the two genotypes.

#### Supplementary Figure S2: Transcriptomic analysis of sensitized hemocytes in Toll$^{10b}$ background.

**A)** Outline of the pipeline followed to generate and analyse the transcriptome of hemocytes from *gcm$^{26}$*, *Toll$^{10b}$* and *Gcm26/+;Toll$^{10b}$/+* larvae. Three replicates were sequenced per genotype.

**B)** Number of reads and percentage of uniquely mapped reads for the 9 transcriptomes.
**C)** GO term enrichment analysis for genes down regulated or up regulated in *Gcm26/+;Toll$^{10b}$/+* compared to *gcm$^{26}$*. The size of the dots represents the enrichment and the color gradient from white to purple shows the significance of the enrichment, expressed in -Log10(p-value) with white being non-significant and blue highly significant.
**D)** Transcriptome comparison of hemocytes from WL3 *gcm$^{26}$* and *Gcm26/+;Toll$^{10b}$/+*. The x-axis represents the average genes expression levels (n=3) and the y-axis the Log2 Fold Change *Gcm26/+;Toll$^{10b}$/+/gcm$^{26}$*. The red dots highlight the genes of the Jak/Stat pathway.
**E)** GO term enrichment analysis for genes down regulated or up regulated in *Gcm26/+;Toll$^{10b}$/+* compared to *Toll$^{10b}$*. The dotplot is represented as in **(C)**. Note that the p-values are higher than in **C**.
**F)** Schematic representation of the Toll signaling pathway. The ligand Spätzle (Spz) binds to the receptor Toll (Tll) and induces the phosphorylation of the inhibitor of NfKB Cactus (cact) through the interaction with the adaptor proteins Myd88 and Tube (Tub) and the tyrosine kinase Pelle (Pll). The phosphorylation of Cact induces its degradation, which release the NFKB transcription factors Dorsal (Dl) and Dorsal-related immunity factor (Dif). Dl/Dif translocate to the nucleus and induce the expression of effector genes such as the antimicrobial peptides *Drosocin* (*Dro*), *Diptericin B* (*DptB*), *Attacin C* (*AttC*) or the serine protease *SPH93* (32).

#### Supplementary Figure S3: *nAchRalpha6* is expressed in resident hemocytes.

**A-C''')** Filet **(A-B'')** and hemocytes **(C-C''')** immunolabellings of L3 larvae *nAchRalpha6-T2A-Gal4/srp(hemo)3xmcherry;UAS-encGFP/+*. The field in **(A-B'')** focus on resident hemocytes located in the dorsal-posterior area of the larva. The hemocytes are labelled with anti-RFP (in red) and the cells expressing *nAchRalpha6-T2A-Gal4* with anti-GFP (in green). In **(A-B'')**, the gray channel represents anti-HRP labelling that shows neurons and in **(C-C''')** it represents anti-P1 labelling that show hemocytes. The complete stack projections are shown in **(A,C-C''')**, a single section is shown in **(B-B'')**.
**D-E''')** Immunolabelling of L3 filets expressing the fusion protein nAchRalpha6-YFP from the endogenous locus of *nAchRalpha6* (45). The YFP is labelled with anti-GFP (in green), the neurons with anti-HRP (in red) and the hemocytes with anti-He (in gray). A single section of a field covering resident hemocytes located in the dorsal-posterior part of the larva is shown in **(D-D''')** and full stack of a field located in the ventral part is shown in **(E-E''')**.
In all fields, the arrowheads indicate the hemocytes expressing nAchRalpha6.

#### Supplementary Figure S4: Quantification of apoptotic hemocytes in nAchRalpha6 knock down and gain of function animals.

**A-B")** Immunolabelling of hemocytes from WL3 *HmldeltaGal4,Dbgtrace* with pH3 mitotic marker **(A-A'')** or with Dcp-1 apoptotic marker **(B-B")**. The hemocytes are labelled with anti-GFP (in green), anti-pH3 (in red) or anti-Dcp-1 (in grey) and the nuclei with DAPI ( in blue). Scale bar: 20 µm.

**C,D)** Quantification of the apoptotic hemocytes in WL3 (120 hAEL) *HmldeltaGal4,Dbgtrace* and *HmldeltaGal4,Dbgtrace,nAchRalpha6-KD* **(A)** or *Pxn>nAchRalpha6 CRISPR GOF* and control **(D)**; n = 12, with more than 300 hemocytes scored for each replicate, p-values were estimated by one-factor ANOVA.

#### Supplementary Table S1: expression matrix from the RNAseq data on sensitized hemocytes in $Toll^{10b}$ background.

Columns A,B: Gene Id, FBgn and Symbol.

Columns C-K: raw read counts for each sample (3 samples *gcm*$^{26}$*/+*, 3 samples *gcm26/+;Toll*$^{10b}$*/+* and 3 samples *Toll*$^{10b}$*/+*.

Columns L-T: normalized read count from DEseq2 on each sample.

Column U: gene symbol

Column V: gene expression base mean across the whole dataset

Columns W-Y: mean expression in *gcm*$^{26}$*/+*, *gcm26/+;Toll*$^{10b}$*/+* and *Toll*$^{10b}$ hemocytes.

Columns Z-AC: Fold change, log2fold change, p-value and adjusted p-value estimated with DEseq2 for the comparison of *Toll*$^{10b}$*/+* and *gcm26/+;Toll*$^{10b}$*/+* hemocytes.

Columns AD-AG: Fold change, log2fold change, p-value and adjusted p-value estimated with DEseq2 for the comparison of *gcm*$^{26}$*/+* and *gcm26/+;Toll*$^{10b}$*/+* hemocytes.

Column AH: genes annotated as transmembrane receptors according to Flybase.

#### Supplementary Figure S1

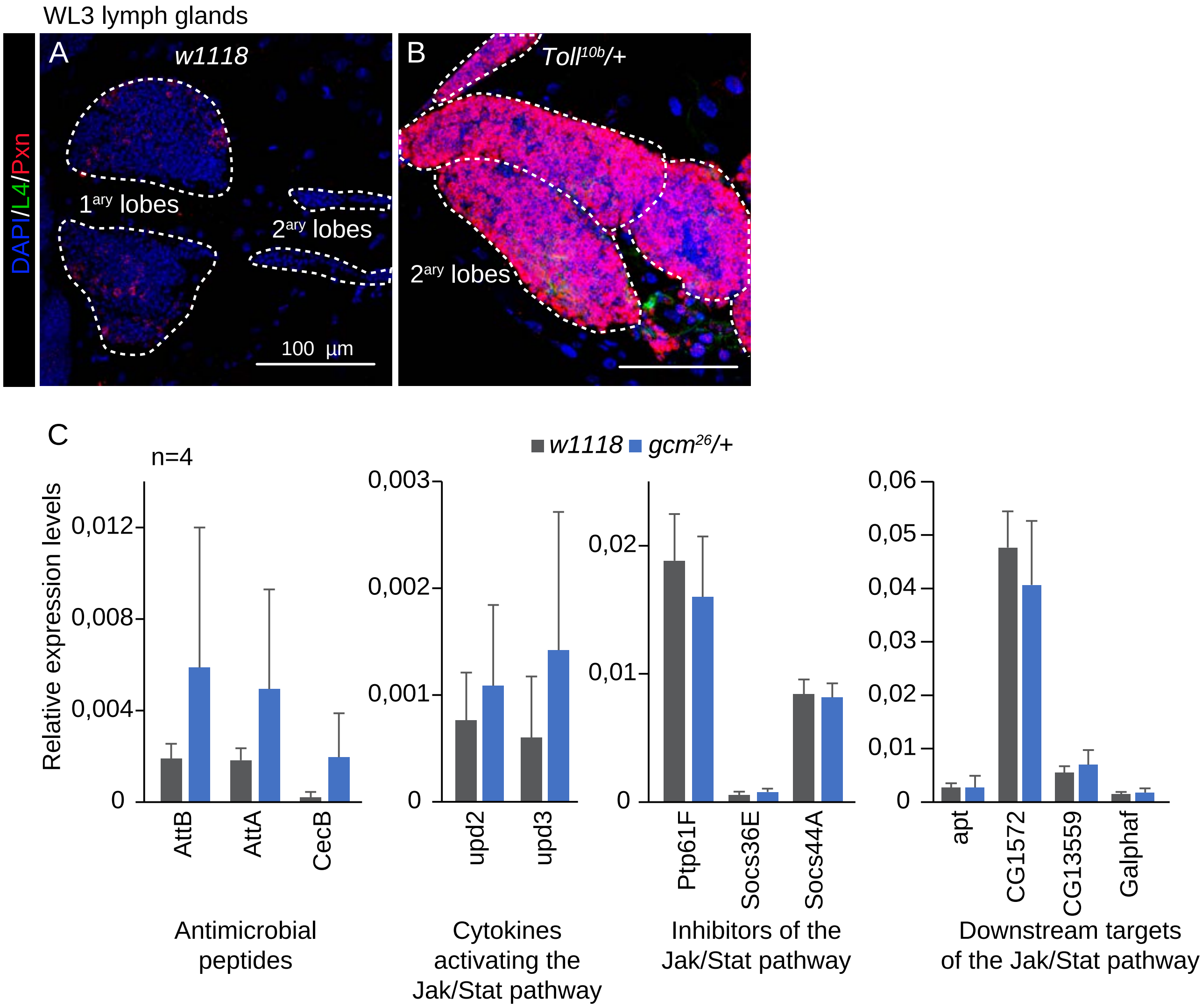

### Supplementary Figure S2

A

| Hemocytes from 100 larvae WL3 (n=3) | Transcriptome generation | Transcriptome analysis |
|---|---|---|
| *gcm^26/+* | Stranded mRNA sequencing | Mapping: Tophat (v1.4.0, genome dm6 ) |
| *Toll^10b/+* | (Illumina Hiseq 2500, | Expression levels: HTseq (v0.6.0) |
| *gcm^26/+;Toll^10B/+* | single end, 50nt) | Differential expression: DESeq2 (v0.99.2) |

B

| Genotype | Replicate | Sample_ID | Number of reads | % of reads uniquely mapped |
|---|---|---|---|---|
| *gcm^26/+* | 1 | WLBZ10 S1 | 33,745,068 | 78.45% |
| *gcm^26/+* | 2 | WLBZ11 S1 | 30,846,975 | 71.87% |
| *gcm^26/+* | 3 | WLBZ12 S1 | 33,718,526 | 76.17% |
| *Toll^10b/+* | 1 | WLBZ1 S1 | 51,491,906 | 86.29% |
| *Toll^10b/+* | 2 | WLBZ2 S1 | 48,991,987 | 86.35% |
| *Toll^10b/+* | 3 | WLBZ3 S1 | 50,968,016 | 86.18% |
| *gcm^26/+;Toll^10B/+* | 1 | WLBZ7 S2 | 62,296,451 | 88.30% |
| *gcm^26/+;Toll^10B/+* | 2 | WLBZ8 S2 | 46,443,939 | 86.64% |
| *gcm^26/+;Toll^10B/+* | 3 | WLBZ9 S2 | 60,029,803 | 87.87% |

C

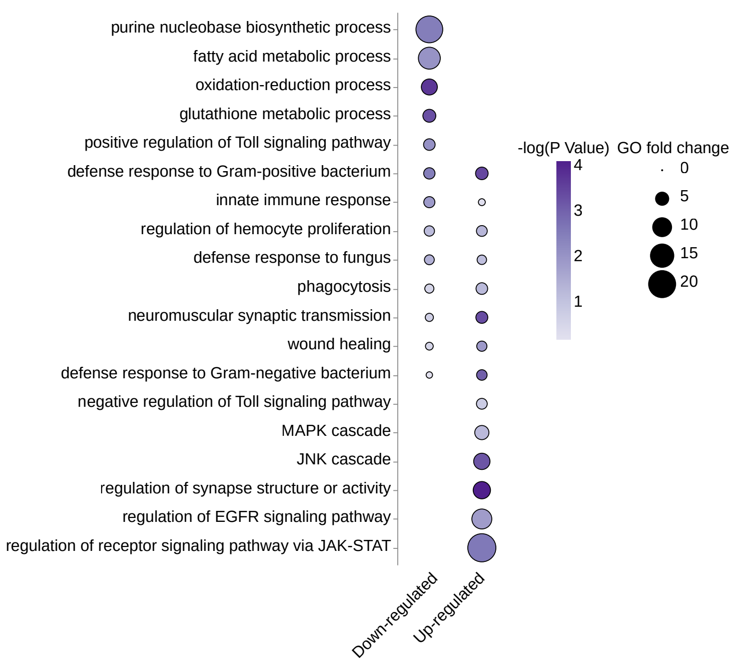


D

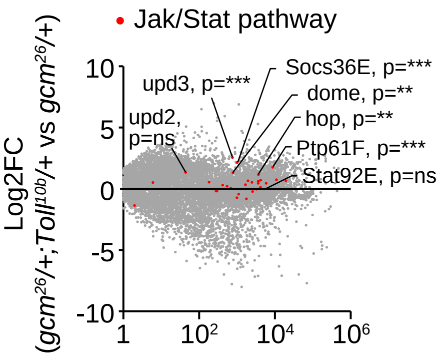


E

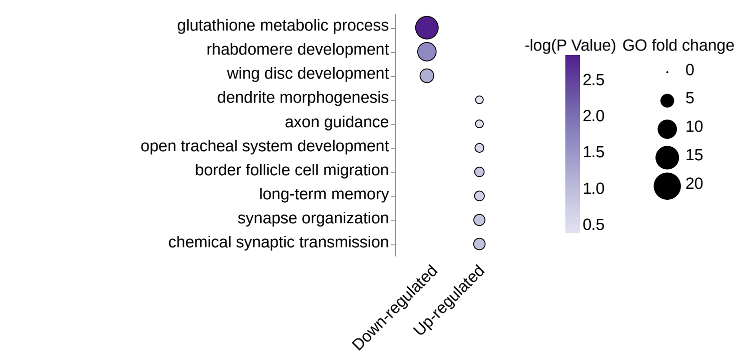


F

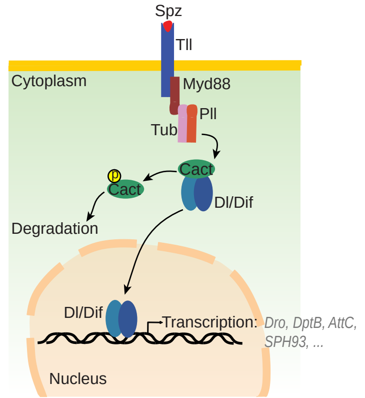

### Supplementary Figure S3

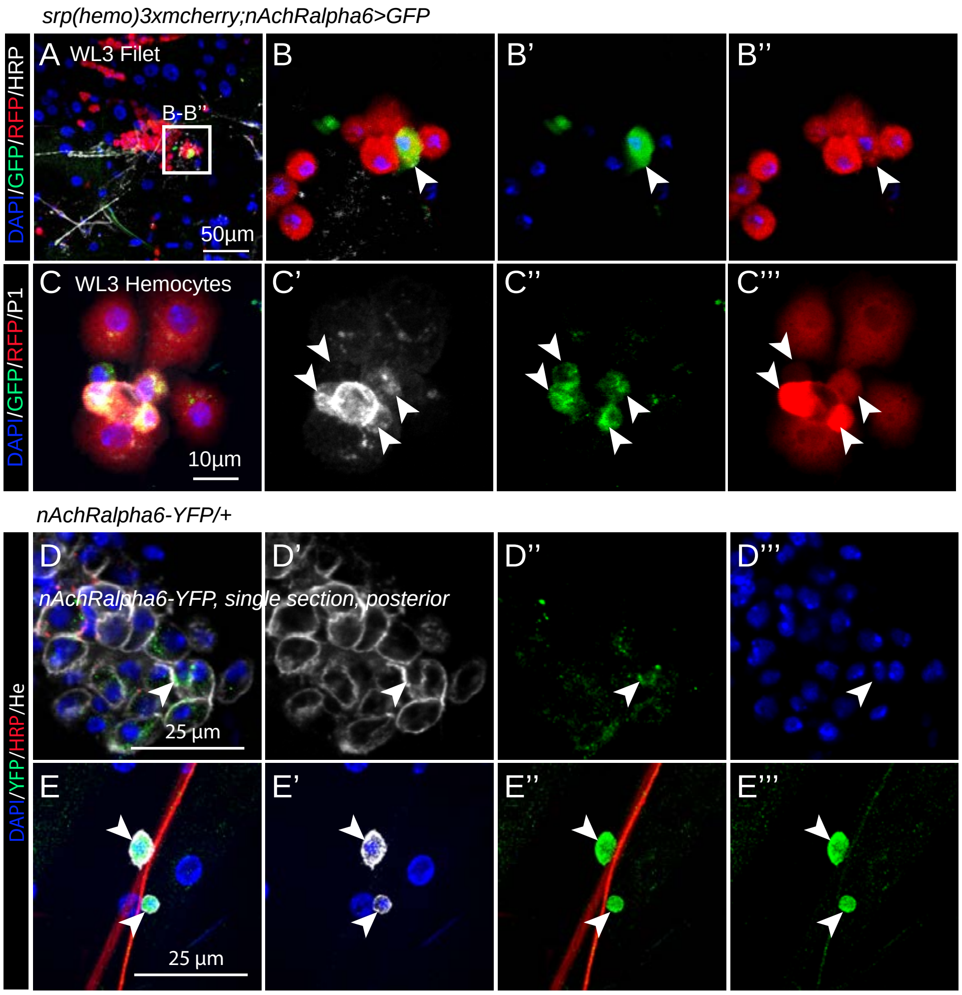

### Supplementary Figure S4

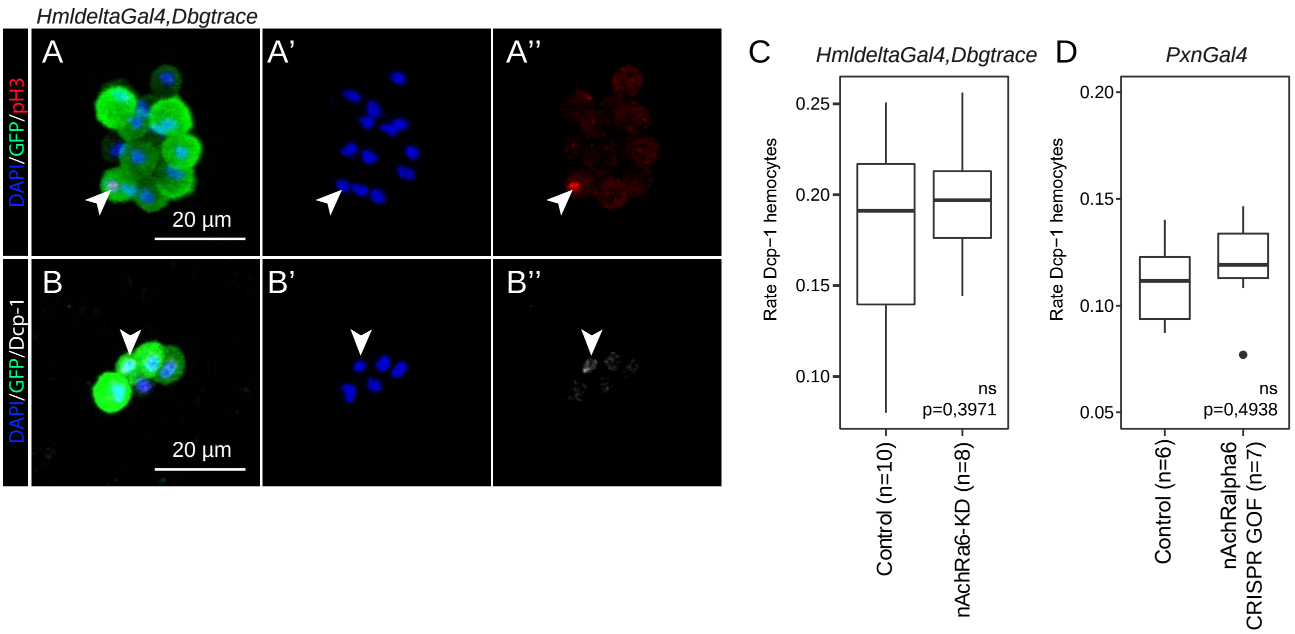

#### Supplementary methods

##### Fly strains and genetics

Flies were raised on standard media at 25°C. The following stocks were used:

| Genotypes | Abbreviation | Origin | Remarks |
|---|---|---|---|
| *gcmGal4/CyO,Tb* | *gcm>* | (Soustelle and Giangrande 2007) | Driver specific to embryonic hemocytes and glia, *gcm* hypomorphic mutation |
| *Toll*$^{10b}$*/TM3_Ser mwh[1] e[1] Tl[*$^{10b}$*]/T(1;3)OR60/TM3. Sb[1] Ser[1]* | *Toll*$^{10b}$*/Ser, TM3* | (Schneider *et al.* 1991; Huang *et al.* 1997) | Constitutively activated Toll. |
| *UAS-gcmRNAi* | *gcm KD* | Bloomington #31519 | dsRNA targeting *gcm* with *UAS* promoter |
| *UAS-gcmF18A* | *gcm GOF* | (Bernardoni *et al.* 1997) | *gcm* gain of function (GOF) under *UAS* promoter |
| *gcm*$^{26}$*/CyOactinGFP* | *gcm*$^{26}$ | (Vincent *et al.* 1996) | Null *gcm* mutation |
| *srp(hemo)Gal4* | *srp(hemo)>* | (Bruckner *et al.* 2004) | Driver specific to embryonic hemocytes |
| *Df(2L)132* | *Df(2L)132* | (Kammerer and Giangrande 2001) | Chromosomal deletion of the locus covering *gcm* and *gcm2*. |
| *srp(hemo)-3xmcherry* | *Srp(hemo) 3xmcherry* | (Gyoergy *et al.* 2018) | Hemocyte reporter |
| *w[*]; P(w[+mC]=UAS-GFP.S65T)eg[T10]* | *UAS-GFP* | Bloomington #1522 | *GFP* reporter under *UAS* promoter |
| *nAchRbeta3-T2A-Gal4* | *nAchRbeta3-T2A-Gal4* | (Kondo *et al.* 2020) | Driver reporting *nAchRbeta3* expression |
| *nAchRalpha6-T2A-Gal4* | *nAchRalpha6-T2A-Gal4* | (Kondo *et al.* 2020) | Driver reporting *nAchRalpha6* expression |
| *Dop1R1-T2A-Gal4* | *Dop1R1-T2A-Gal4* | (Kondo *et al.* 2020) | Driver reporting *Dop1R1* expression |
| *w[1118]; P(w[+mC]=Hml-Gal4.Delta)2* | *HmldeltaGal4* | Bloomington #30139 | Driver specific to hemocytes |
| *PxnGal4 (chr3)* | *PxnGal4* | Gift from Pr. D. Hultmark | Driver specific to hemocytes |
| *y[1]v[1];P(y[+t7.7]v[+t1.8]=TRiP.JF01851)attP2* | *UAS-5HT1B-RNAi* | Bloomington #25833 | dsRNA targeting *5-HT1B* with *UAS* promoter |
| *y[1]v[1];P(y[+t7.7]v[+t1.8]=TRiP.HM04077)attP2* | *UAS-Dop1R1-RNAi* | Bloomington #31765 | dsRNA targeting *Dop1R1* with *UAS* promoter |
| *y[1]v[1];P(y[+t7.7]v[+t1.8]=TRiP.JF03126)attP2* | *UAS-Ekar-RNAi* | Bloomington #28506 | dsRNA targeting *Ekar* with *UAS* promoter |
| *y[1]v[1];P(y[+t7.7]v[+t1.8]=TRiP.JF01853)attP2* | *UAS-nAchRalpha6-RNAi* | Bloomington #25835 | dsRNA targeting *nAchRalpha6* with *UAS* promoter |
| *UAS-nAchRbeta3-RNAi (chr2)* | *UAS-nAchRbeta3-RNAi* | VDRC ID: KK 101868 | dsRNA targeting *nAchRbeta3* with *UAS* promoter |

| | | | |
|---|---|---|---|
| *y[1]v[1];P(y[+t7.7]v[+t1.8]=TRiP.HMS05847)attP40* | UAS-RFP-RNAi | Bloomington #67984 | dsRNA targeting *RFP* with *UAS* promoter (Control) |
| *nAchRalpha6-YFP* | *nAchRalpha6-YFP* | (KORONA *et al.* 2021) | Reporter of *nAchRalpha6* |
| *nAChRalpha6[DAS1]* | *nAChRα6*$^{DAS1}$ | Bloomington #9685 | Null mutation of *nAchRalpha6*. The genetic background was homogenised by chromosome replacement with the wild type stock Oregon-R |
| *nAChRalpha6[DAS2]* | *nAChRα6*$^{DAS2}$ | Bloomington #9686 | Null mutation of *nAchRalpha6*. The genetic background was homogenised by chromosome replacement with the wild type stock Oregon-R |
| *w[*];P(w[+mC]=UAS-FLP.Exel)3,P(w[+mC]=Ubi-p63E(FRT.STOP)Stinger)15F2* | *gtrace 3* | Bloomington #28282 | Lineage tracing stock on chr 3 |
| *y,w,UAS-FLP;Act5C-FRT-y+-FRT-GAL4,UAS-GFP* | *gtrace 2* | (HONTI *et al.* 2010) | Lineage tracing stock on chr X and 2 |
| *y,w,UAS-FLP;Act5C-FRT-y+-FRT-GAL4,UAS-GFP ;P(w[+mC]=UAS-FLP.Exel)3,P(w[+mC]=Ubi-p63E(FRT.STOP)Stinger)15F2* | *Dbgtrace* | Build in this study | Stock carrying the two lineage tracing constructs *gtrace 2* and *gtrace 3*. |
| *y[1]sc[*]v[1]sev[21];P(y[+t7.7]v[+t1.8]=TRiP.HMC05021)attP40* | *UAS-ChAT-RNAi* | Bloomington #60028 | dsRNA targeting *ChAT* under *UAS* promoter |
| *y[1] w[*]; P(w[+m*]=nSyb-GAL4.S)3* | *nSyb-Gal4* | Bloomington #51635 | Neuronal *Gal4* driver |
| *TI(2A-lexA::GAD)ChAT[2A-lexA]* | *ChAT-T2A-LexA* | Bloomington #84379 | Cholinergic neuron *LexA* driver |
| *w[1118]; P(w[+mC]=ChAT-GAL4.7.4)19B/CyO, P(ry[+t7.2]=sevRas1.V12)FK1* | *ChAT-Gal4* | Bloomington #6798 | Cholinergic neuron *Gal4* driver |
| *21-7-Gal4* | *21-7-Gal4* | (HU *et al.* 2017) | Multidendritic neuron *Gal4* driver |
| *y[1]w[*];P(w[+mW.hs]=GawB)109(2)80,P(w[+mC]=UAS-mCD8::GFP.L)LL5 (1;2)* | *109(2)80-Gal4* | Bloomington #8768 | Multidendritic neuron *Gal4* driver |
| *P(w[+mW.hs]=GawB)OK6 (2)* | *OK6-Gal4* | Bloomington #64199 | Motorneuron *Gal4* driver |
| *w*; P(Tdc2-GAL4.C)2* | *Tdc2-Gal4* | Bloomington #9313 | Octopaminergic neuron *Gal4* driver |
| *w[1118]* | *w1118* | | Wild type |
| *Oregon-R* | *Oregon-R* | | Wild type |
| *w[1118]; P(y[+t7.7] w[+mC]=UAS-Cas9.C)attP2* | *UAS-Cas9* | Bloomington #54595 | |

| | | | |
|---|---|---|---|
| *y[1] sc[*] v[1] sev[21]; P(y[+t7.7] v[+t1.8]=TKO.GS00614)attP40* | *nAchRalpha6-TKO* | Bloomington # 76441 | Expresses sgRNAs ubiquitously targeting *nAChRalpha6* for knock down with *Cas9* |
| *y[1] w[*]; P(y[+t7.7] w[+mC]=UAS-3xFLAG.dCas9.VPR)attP40* | *UAS-dCas9-VPR* | Bloomington #66561 | |
| *y[1] sc[*] v[1] sev[21]; P(y[+t7.7] v[+t1.8]=TOE.GS01969)attP40* | *nAchRalpha6-TOE* | Bloomington #80510 | Expresses sgRNAs ubiquitously targeting *nAChRalpha6* for overexpression with *dCas9-VPR* |
| *nAchRalpha6-TKO ;PxnGal4* | | Build in this study | Crossed with *UAS-Cas9* to generate *nAchRalpha6 CRISPR LOF* |
| *nAchRalpha6-TOE ;PxnGal4* | | Build in this study | Crossed with *UAS- dCas9-VPR* to generate *nAchRalpha6 CRISPR GOF* |
| *nAchRalpha6-TKO/UAS-Cas9;PxnGal4/+* | *nAchRalpha6 CRISPR LOF* | Build in this study | |
| *UAS-Cas9/+;PxnGal4/+* | *Control nAchRalpha6 CRISPR LOF* | Build in this study | |
| *nAchRalpha6-TOE/UAS-dCas9-VPR;PxnGal4/+* | *nAchRalpha6 CRISPR GOF* | Build in this study | |
| *UAS- dCas9-VPR/+;PxnGal4/+* | *Control nAchRalpha6 CRISPR GOF* | Build in this study | |

#### Estimation of the penetrance of the melanotic tumors

Overnight lays of the parental crosses were incubated at 25°C until the wandering 3rd instar larval (WL3) stage. The larvae were then washed with water and tumor penetrance was determined by assessing the percentage of larvae carrying one or more tumors. More than 50 larvae were scored per genotype. The p-values were estimated using the chi-squared test for frequency comparisons.

#### Hemocyte counting

Parental crosses were allowed to lay eggs on collecting plates for 3hrs at 25°C. After 24hrs at 25°C, 100 1st instar larvae were transferred from the collecting plate to fresh vials (to normalise the number of animals per vial, avoid overcrowding and stage the larvae). At wandering stage (~116hrs after egg laying (hAEL)), 10 female larvae were washed and bled in PBS containing few crystals of N-phenylthiourea ≥98% (PTU) (Sigma-Aldrich P7629) to prevent hemocyte melanisation (LERNER AND FITZPATRICK 1950). During the bleeding procedure, the cuticle was scraped with forceps to recover the sessile hemocytes (PETRAKI *et al.* 2015). The hemocytes were then numerated on a Malassez counting cell and reported as number of hemocytes per larva. At least 3 replicates were performed for each genotype. The counts were compared using ANOVA followed by post hoc Tuckey HSD or bilateral student test as mentioned on each graph.

For the lamellocytes, 10 WL3 were treated as stated above and bled in PBS/PTU. The hemocytes were transferred onto a slide using the Cyto-Tek® 4325 Centrifuge (Miles Scientific). Samples were then fixed for 10min in 4% paraformaldehyde/PBS at room temperature (RT), incubated with blocking reagent (Roche) for 1hr at RT, incubated overnight at 4°C with primary antibodies targeting hemocyte (rabbit anti-Serpent, (TREBUCHET *et al.* 2019))and lamellocytes (Mouse anti-L4 kindly provided by I. Ando (HONTI *et al.* 2010)) diluted in blocking reagent. The slides were then washed three times for 10min with PTX (PBS, 0.3% triton-x100), incubated for 1hr with secondary antibodies, washed twice for 10min with PTX, incubated for 20min with DAPI to label nuclei (Sigma-Aldrich) (diluted to $10^{-3}$ g/L in blocking reagent), and then mounted in Vectashield® (Vector Laboratories). The slides were analysed by confocal microscopy. Each immunolabelling was carried out on three independent trials. The p-values were estimated after variance analysis using bilateral student test.

#### Lymph gland, filet and whole larva immunolabelling

Lymph glands of the indicated stages were dissected in Ringer's solution (pH 7.3-7.4), fixed for 30min in 4% PFA/PBS at 4°C, washed three times for 10min with PTX, incubated with blocking reagent for 1hr at 4°C, incubated overnight at 4°C with primary antibodies, washed three times for 10min with PTX, incubated for 1hr with secondary antibodies, washed two times for 10min with PTX, incubated for 20min with DAPI and then mounted on slides in Vectashield®. The slides were analysed by confocal microscopy.

The filets were prepared in cold PBS following the protocol from Brent et al. with modifications (Brent *et al.* 2009). Briefly, to expose resident hemocytes located in dorsal and lateral positions, the larvae were dissected ventrally or laterally on a cold block to prevent dislodging the resident hemocytes. The organs were removed gently from the pinned larvae. Then the filets were fixed in 4% PFA in 1x PBS for 30min and treated as described above for the lymph glands.

Whole larva labelling was carried out following the protocol from (Manning and Doe 2017) with modifications. Briefly, the feeding L3 were collected and washed with water, treated 10min with 100 % bleach and rinsed in water. Then the larvae were treated with proteinases for 1h at 37°C in CCD buffer (chitinase (5 U/ml), chymotrypsin (100 U/ml) in HEPES buffer (25 mM) with 1,5% (vol/vol) DMSO) supplemented with 7 M urea to improve antibody penetration. The larvae were then fixed in 9% (vol/vol) PFA in PBS with heptane for 30 min at room temperature and treated with heptane and methanol (cracking procedure). A second fixation was carried out for 7 days at -20°C with 2% PFA in methanol. Heptane was added to increase the permeability of the animals for the immunostaining. Following this, the larvae were treated with Proteinase K to increases the coloration in deep tissue, blocked overnight with blocking reagent (Roche) in PTX, incubated 7 days with the primary antibodies and 1 day with the secondary antibodies.

For all the immunolabellings, the following primary antibodies were used: mouse anti-Hemese (1/30), (Kurucz *et al.* 2003) and mouse anti-P1 (1/30), kindly provided by I. Ando (Kurucz *et al.* 2007); rabbit anti-PH3 (1/1000) (Upstate biotechnology #06-570); chicken anti-GFP (1/500) (abcam #13970); rat anti-RFP (1/500) (Chromotek #5F8-100); rabbit anti-HRP (1/500) to label neurons (Jan and Jan 1982); mouse anti-Dlg (DSHB Cat# 4F3 anti-discs large, RRID:AB_528203); mouse anti-Brp (DSHB Cat# nc82, RRID:AB_2314866) and the following secondary antibodies: Cy3 Donkey Anti-Mouse IgG (Jackson ImmunoResearch Labs Cat# 715-165-151, RRID:AB_2315777); Cy3 Goat Anti-Rat IgG (Jackson ImmunoResearch Labs Cat# 112-165-167, RRID:AB_2338251); Cy5 Goat Anti-Mouse IgG (Jackson ImmunoResearch Labs Cat# 115-177-003, RRID:AB_2338719); Cy5-AffiniPure Goat Anti-Rat IgG (H+L) (Jackson ImmunoResearch Labs Cat# 112-175-167, RRID:AB_2338264 ) and FITC anti-chicken (Jackson ImmunoResearch Labs Cat#703-095-155).

#### Confocal imaging

Leica SP8 inverted-based microscope equipped with 20, 40 and 63X objectives was used to obtain confocal images. GFP/FITC was excited at 488nm; the emission filters 498-551 were used to collect the signal. Cy3 was excited at 568nm; emission filters 648-701 were used to collect the signal, and Cy5 was excited at 633nm; emission signal was collected at 729-800nm. A step size between 0.2 and 2µm was used to collect the Z-series of images, which were then treated with Fiji (Schindelin *et al.* 2012). For each set of images, the intensity of the signals was set to the same threshold in order to compare the different genotypes.

#### Stranded RNA sequencing on hemocytes from WL3 larvae

For each replicate, the hemocytes were bled from 100 WL3 of the following genotypes: (1) *gcm*$^{26}$*/+* issued from the cross *gcm*$^{26}$*/CyOGFP* x *w*$^{1118}$, (2) *Toll*$^{10b}$*/+* issued from the cross *Toll*$^{10b}$*/Ser,TM3* x *w*$^{1118}$ and (3) *gcm*$^{26}$*/+;Toll*$^{10b}$*/+* issued from the cross *gcm*$^{26}$*/CyOGFP* x *Toll*$^{10b}$*/Ser,TM3*. The RNA were extracted using Tri reagent (Sigma) following the manufacturer's instruction. Library preparation and sequencing were carried out by the genomeast facility (https://www.igbmc.fr/plateformes-technologiques/genomeast) at the IGBMC. Stranded mRNA libraries were prepared with Directional mRNA-Seq (SamplePrep). The libraries were sequenced on the Illumina Hiseq 2500 as single-end 50 base reads following Illumina's instructions. Image analysis and base calling were performed using RTA 1.18.61 and CASAVA 1.8.2. The reads were mapped to the *Drosophila* genome dm6 using Tophat (v1.4.0) (Trapnell *et al.* 2009) with the following parameters: library type FR second strand, anchor length = 8, minimum intron length = 70, maximum intron length = 500000. The bam files were then converted to count files using Htseq count (Anders *et al.* 2015) on stranded data and compared using DESeq2 (Varet *et al.* 2016). The graphical representations were built using R version 4.1.0, R studio version 1.4.1717 and the packages ggplot2 (Wickham 2016) and pheatmap (RRID:SCR_016418).

The RNA-seq data have been deposited in the ArrayExpress database at EMBL-EBI (www.ebi.ac.uk/arrayexpress) under accession number ***E-MTAB-11970.***

#### GO term enrichment analysis.

GO term enrichment analysis was done using PANGEA for multiple lists (Hu *et al.* 2023) using the EXP GO Biological Processes database. Genes presenting a base mean expression above 100, an upregulation above 2 fold (Log2 fold change > 1) and an adjusted p-value below 0.01 were selected for the analysis.

#### RNA extraction and qPCR

For the qPCR validation of the transcriptomic data, 20 WL3 of the indicated genotypes were bled on ice cold PBS. The cells were then centrifuged at 1200 rpm at 4°C and RNA isolation was performed with Tri-reagent (Sigma) following the manufacture's protocol. The DNase treatment was done with the TURBO DNA-free kit (Invitrogen) and the reverse transcription (RT) with the Super-Script IV (Invitrogen) using random primers. The qPCR assays were done with FastStart Essential DNA Green Master (Roche) with the primers are listed below.

| Target | Forward primer | Reverse primer |
|---|---|---|
| *AttA* | GGCCCATGCCAATTTATTCA | AGCAAAGACCTTGGCATCCA |
| *CecA2* | CATTCTGGCCATCACCATTGGACA | GTGTGCTGACCAACACGTTCGATT |
| *CecB* | TTCGTCTTTGTGGCACTCATCCTG | GGTATGCTGACCAATGCGTTCGAT |
| *GstE7* | CGATGATTCCCAAGGAGCGT | GGTCGTGTCCACCTTTACGA |
| *GstE1* | CCGTACACCAGGGTCTGAAG | ATTGAGGCGATCCAACCAGG |
| *nAchRalpha6* | AGAGCCACGCGATACAAACA | GGTGGACAGCAGATGGTTCA |
| *Rp49* | GACGCTTCAAGGGACAGTATCTG | AAACGCGGTTCTGCATGAG |
| *Act5C* | GCAGCAACTTCTTCGTCACA | CTTAGCTCAGCCTCGCCACT |

The expression levels were calculated relative to the two housekeeping genes Rp49 and Act5C levels using the ΔCt formula: $2^\wedge(\text{average}(Ct_{Rp49}, Ct_{Act5c}) – Ct_{target})$. Triplicate were done for each genotype and the levels were compared using bilateral student test after variance analysis.

#### DHE, pH3 and Dcp-1 quantification

For the estimation of the ROS levels, single *Toll*$^{10b}$*/+* and *gcm*$^{26}$*/+;Toll*$^{10b}$*/+* WL3 were bled in 20 µL of Schneider medium containing PTU on a 12 wells chamber slide (1 larva per well). The hemocytes were left to decant for 20min and incubated in Schneider medium containing 10 µM dihydroethidium (DHE, Sigma) and Hoechst for 30min at RT in a wet chamber. The endogenous ROS convert the DHE to the fluorescent 2-hydroxyethidium (ZHAO *et al.* 2005), which absorbs 488nm light and emits photons at ~600nm. Hoechst labels the nuclei of the hemocytes. Following this, the medium was replaced by fresh Schneider medium and the slides were scanned using CellInsight CX7 instrument (Cellomics) at 20x magnification (25 fields per well). The intensity of the DHE within the nuclei delimited by the Hoechst signal was estimated using HCS Studio software and Colocalisation Bioapplication, for at least 7 larvae per genotype with more than 200 cells per larva. The DHE intensity averages were compared using bilateral student test.

For the estimation of the number of mitotic and apoptotic hemocytes, single WL3 were bled in 20 µL of Schneider medium containing PTU on a 12 wells chamber slide (1 larva per well). The hemocytes were left to decant for 20 min, fixed for 10min with 4% PFA in 1x PBS, washed with PTX, incubated for at least 1hr with blocking reagent and labelled with primary antibodies targeting the mitotic marker pH3 or the apoptotic marker cleaved Dcp-1 and secondary antibody coupled with Cy5, as well as DAPI to label the nuclei. The images were acquired using CellInsight CX7 instrument (Cellomics) at 20x magnification (100 fields per well). The image analysis was done using HCS Studio software and Colocalisation Bioapplication. The region of interest was defined using DAPI labelling and the pH3 or Dcp-1 intensity was measured for each region. The cells displaying a pH3 intensity above 6 times the median intensity or a Dcp-1 intensity above 1.8 times the median intensity of the well (determined empirically over 3 wells, **Supplementary Figure S4A,B**) were considered as proliferative for pH3 or apoptotic for Dcp-1. The rates of proliferation were compared using ANOVA.
